# An Immune Cell Recirculation-Enabled Microfluidic Array to Study Dynamic Immunotherapeutic Activity in Recapitulated Tumor Microenvironment

**DOI:** 10.1101/2022.09.09.507299

**Authors:** Chun-Wei Chi, Yeh-Hsing Lao, AH Rezwanuddin Ahmed, Siyu He, Taha Merghoub, Kam W. Leong, Sihong Wang

## Abstract

The efficacy of immunotherapeutic treatment protocols to enable immune cell mediated treatment of cancer is significantly modulated in the presence of tumor microenvironment (TME) which is a key factor in providing both a physical barrier and immunosuppressive stimuli. Herein, we developed a recirculating, high-throughput microfluidic cell array to capture these crucial players – cytotoxic T cells in circulation, endothelium, and tumor stroma. The system consisted of a three-layered cell array spatially emulating TME, with T cell circulation sustained via fluidic recirculating circuits. This allowed us to study the dynamic TME/circulation system and cancer cell response thereof. The system further revealed that tumor endothelium exhibited a hindrance to T cell infiltration into the breast cancer tumor compartment, which was alleviated when treated with anti-human PD-L1 antibody. The other key stromal component, cancer associated fibroblasts, further attenuated T cell infiltration, and led to reduced apoptosis activity in cancer cells. These results confirm the capability of our tumor-on-a-chip system to recapitulate some key immune cell interactions with the reconstructed TME, along with demonstrating as the feasibility of using this system for high-throughput cancer immunotherapeutic screening.

---

Immunotherapeutics, including immune checkpoint inhibitors and chimeric antigen receptor T cell (CAR-T cell) therapy, have emerged in the past decade. Their success in hematologic cancers have led to more than 4,800 active clinical trials to target nearly 50 other cancer types, indicating that a massive race in translation of immunotherapeutics has been taking place.^1, 2^ Despite some success, application of immunotherapeutics for treating solid tumors have mixed outcomes in clinical trials, wherein many of them work only in a small portion of the patients,^3–5^ hinting at the possibility of undiscovered factors during preclinical validations. Particularly, the interplay between immunotherapeutics, tumor cells and immune cells, such as T cells, could mediate treatment efficacy. For example, high expression of the programmed cell death ligand 1 (PD-L1) on the cancer cell surface could negatively affect T cell infiltration, while blocking of surface PD-L1 by antibodies may improve T cell infiltration into the solid tumor microenvironment (TME). Interactions with the stromal cells in TME could also lead to different drug responses.

Therefore, to screen immunotherapeutics at a clinically relevant caliber, a proper model that emulates the cell-cell interaction and cell-drug modulation is therefore needed. Although murine models are the preclinical gold standards for drug screening, the intrinsic differences between mouse and human immune systems can present irreconcilable differences in treatment outcomes indicated from screening.^6–12^ In contrast, tumor-on-a-chip enables *in vitro* screening with physiologically mimicked TME and exemplifies some characteristics closer to the clinical scenario, which could be missing in non-human *in vivo* or xenograft models. Our group and others have reported different tumor chips with the capabilities of screening cancer drugs in a high-throughput manner or studying cell-cell interactions. Compared with conventional drug screening, tumor chips maintaining 3D culture supported by media flow better mimics the *in vivo* morphological and functional characteristics.^13–16^ A few new designs have been recently developed to investigate immune cell migration, where immune cells were either embedded in gel with cancer cells or seeded onto the gel surface.^17–21^ Although direct contact between stationarily placed cancer and immune cells allow to study some interactive behavior, several key cellular dynamics (e.g., immune cell circulation and transmigration) cannot be seen in these implementations, which leads to predictive drug screening results of high inaccuracy when translated to a physiological milieu.

To address these, here we develop a recirculating high-throughput cell array system that allows to study the dynamic interaction between immunotherapeutic agents, tumor and T cells (**Figure 1A**). The system amalgamates two major components– fluidic recirculating circuits and microfluidic cell array (μFCA). Each recirculating circuit provides unidirectional flow without the traditional use of peristaltic pumps, which maintains flow-guided interactions between T cells and the endothelial layer, while avoiding some undesired, pump-related mechanical stresses on T cells.^22–24^ The μFCA reconstructs the spatial configuration of microvessel and TME, permitting interactions between tumor, stroma, and T cells. Our system is therefore capable of studying the interplay between four key physiological tumor components – T cells, tumor cells, microvasculature, and the remaining tumor stroma - modulated in the presence of immunotherapeutic agents. Recapitulating these dynamic interactions can present a more faithful representation of physiological outcomes.

**Figure 1.**
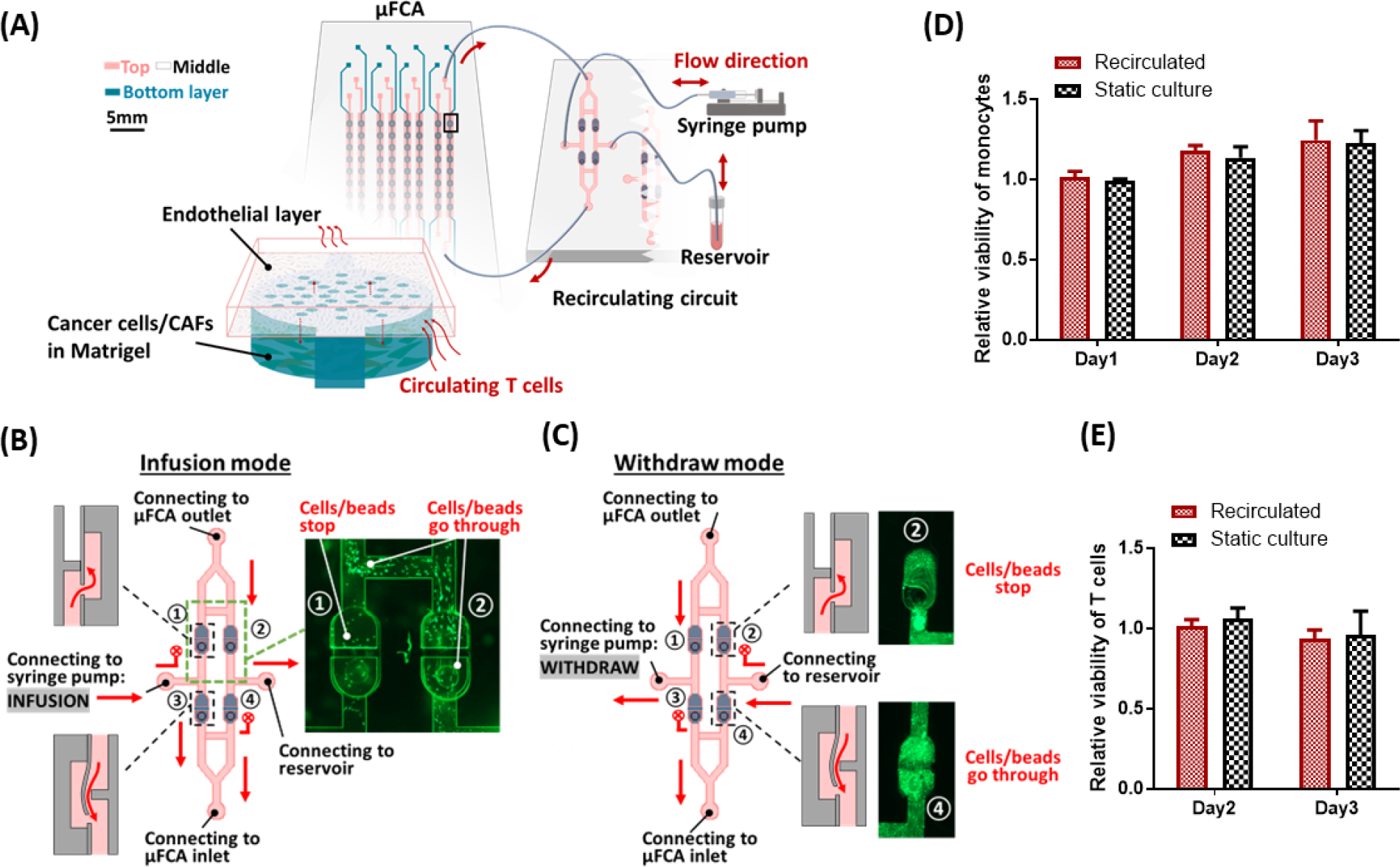
Design and characterization of our recirculating tumor-chip system. (A) Schematic illustration of our setup of microfluidic cell array (μFCA), the recirculating circuit, reservoir, and a syringe pump with an enlarged unit of μFCA. Operation of our recirculating circuit when the connected syringe pump set in (B) “infusion” mode and (C) “withdraw” mode (insert in (B) shows calcein-stained mouse CD8^+^ T cells following the flow direction and insert (C) shows fluorescent beads stopped and bypass the check valves). Viability of (D) human monocytes and (E) mouse CD8^+^ T cells after recirculating in the system. Results are presented as average ± standard deviation (S.D.; n = 3).

Our recirculating circuit consists of four microfluidic one-way check valves to provide unidirectional flow to the μFCA module, which realizes T cell circulation through simple controls via syringe pump’s infusion and withdraw modes. To validate the design, we first tested it with 2 μm-diameter fluorescent latex beads and then with calcein AM-stained mouse CD8^+^ T cells. By tracing the beads/cells in the circuit, we confirmed that the check valves properly controlled the flow. In the infusion mode, the syringe pump pushed the beads/cells-containing fluid through Check Valve 3, thereby guiding the beads/cells into the μFCA through its inlet. On the other hand, Check Valves 1 and 4 are closed to the infusion direction, thus fluid exiting out from μFCA could be directed to the reservoir through the opened Check Valve 2 (**Figure 1B**). During the withdraw mode, the syringe pump pulled fluid out from the reservoir into μFCA through Check Valve 4, while flow exiting from the μFCA outlet refilled the syringe via Check Valve 1, since Check Valves 2 and 3 remained closed during the withdrawal phase (**Figure 1C**). After confirming the recirculating circuit worked as expected, we checked if our check-valve design had any negative effects on the viability of recirculating immune cells because of any possible, undesired stresses. As this effect could be size dependent, we chose primary mouse CD8^+^ T cell and a relatively larger cell type, human monocyte line (THP-1), for validation. When compared with the viability seen in the static culture condition, we did not see significant reduction in cell viability even for the larger THP-1 cell after a 3-day long recirculation in our system (**Figures 1D** and **E**). These show that our recirculating design is superior to the conventional approach with peristaltic pump, known to generate mechanical stress to the recirculating cells. ^22–24^

In the cell array module, the three-layer structure of each unit is similar to what we reported previously,^16, 25^ where the cancer cells and stromal cells are embedded in Matrigel in the bottom chamber, and the endothelium formed by human microvascular endothelial cells is in the top channel connecting different units. Pores between the bottom chamber and the top channel enable nutrient-waste exchange and cell-cell interactions (**Figure 1A**). **Figure 2A** shows one typical unit in our μFCA under fluorescence microscopy. The RFP-tagged triple negative breast cancer (TNBC) cells (MDA-MB-231) that were seeded with Matrigel subsequently self-formed an aggregate in the bottom chamber, while human microvascular endothelial cells aligned into a monolayer in the top channel. Notably, by reconfiguring the orientation of top channel and bottom chamber from parallel to orthogonal orientation and integrating with pneumatic valves, the throughput of one μFCA could be increased from 8 conditions (with 8 technical repeats) to an array of 8×8 conditions, where top layer channels run as columns and bottom layer channels as rows. (**Figure 2B**). In the bottom channel, we added a normally closed pneumatic valve between two adjacent bottom chambers to control the sample separation, the valve’s opening being controlled by a pneumatic channel sitting above (**Figure 2C**). Application of negative pressure deforms the valve into an open state, allowing cell seeding throughout the bottom channel. After seeding, removal of the negative pressure restores the default closed state of the valve, physically isolating chambers from crosstalk. (**Figures S1A and S1B**, **Supporting Information**). To confirm no crosstalk between groups when the valve was closed, we used the fluorescent dye, sodium fluorescein, to test our valves. We first filled the whole μFCA with water, followed by introducing dye to every alternate column in the top layer after the valves were closed. As a result, sodium fluorescein would diffuse into the bottom chambers under the dye-filed columns. Water was maintained in the remaining columns of the top layer, creating a columnar alternating array of dyed and clear channels. Properly closed valves between chambers should ensure no cross-talk between the clear and dyed columns. Considering the chambers in the bottom layer receiving dye as a source and those receiving water as a sink, the relative intensity of sodium fluorescein in the sink chambers was minimal (<5%), with no significant leakage found in 24 h measurement (**Figure S1C, Supporting Information**). To further validate this design, MDA-MB-231 cells with blue fluorescence nucleus dye were mixed with Matrigel and seeded into every other row (bottom chambers), and Texas Red-labeled 40kDa dextran was then introduced to every other column (top layer). Similarly, as shown in **Figure 2D**, there was no dextran diffusing across the closed valves, confirming no crosstalk between the valve-isolated chambers under the continuous perfusion.

**Figure 2.**
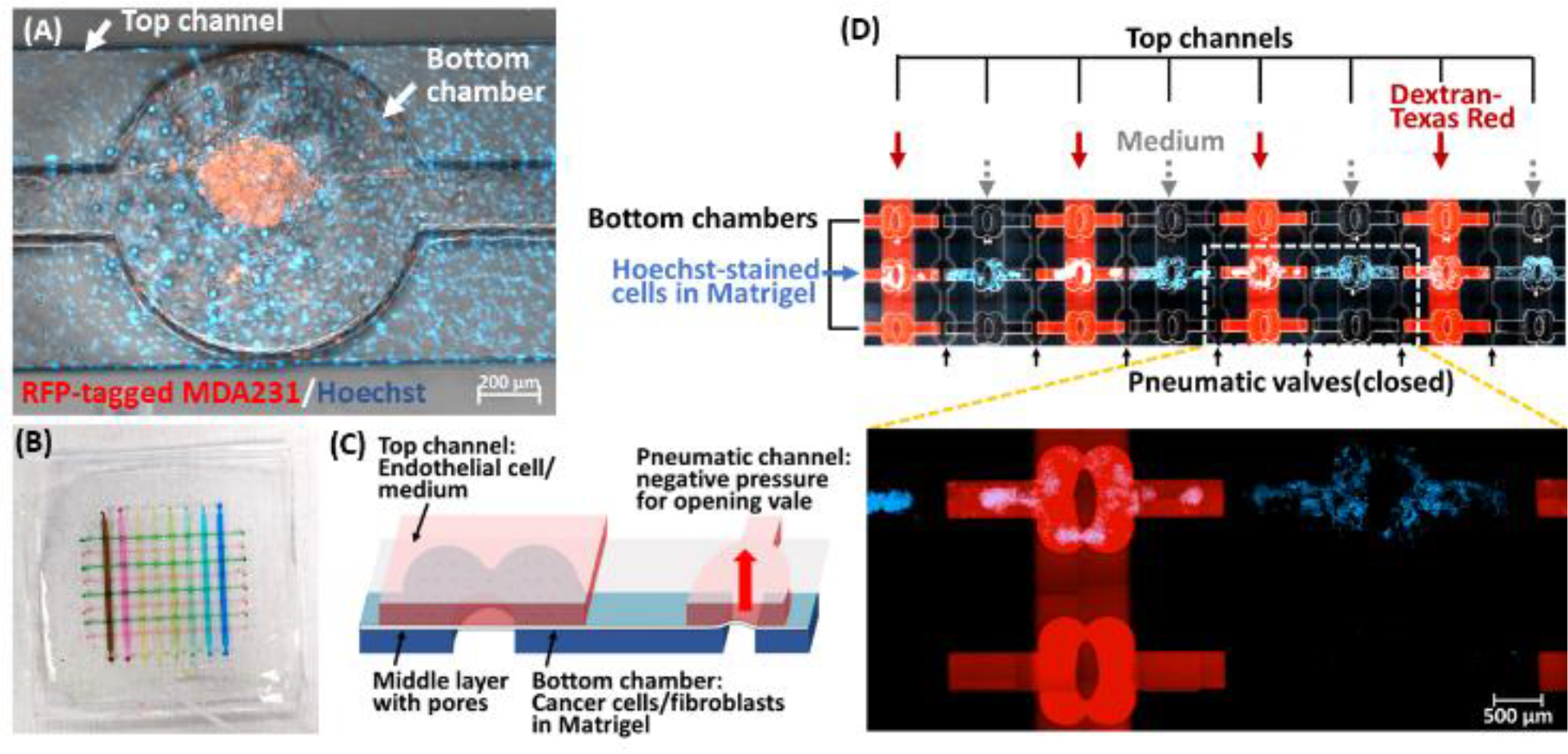
Microfluidic cell array (μFCA) design and the high-throughput configuration. (A) Representative image of one unit in μFCA with the parallel configuration. RFP-tagged MDA-MB-231 cells with Matrigel were seeded in the bottom chamber, while endothelial cells on the top channel formed a monolayer. All the cells were stained with Hoechst-33342 (blue). (B) Representative photo of our high-throughput 8×8 μFCA configuration, consisting of 8 channels and 8 chambers (each channel filed with a unique color dye). (C) Schematic illustration of one unit with the pneumatic valve in the high-throughput 8×8 μFCA configuration with orthogonal layer orientation. (D) Representative images of our high throughput μFCA when validated with cancer cells. MDA-MB-231 cells stained with Hoechst-33342 (blue) were seeded every other row (bottom chambers). The 40kDa Dextran-Texas Red was continuously introduced in every other column (top channels) while the valves were closed.

In order to study the role of T cell dynamics during immunotherapeutic drug responses, we picked the human MHC nonrestricted cytotoxic T-cell line used in clinical trials, TALL-104.^26, 27^ TALL-104 is derived from a 3-year-old patient with acute lymphocytic leukemia and expresses typical surface markers of both cytotoxic T lymphocyte and natural killer cell, including CD3^+^/TCRαβ^+^, CD8^+^, CD56^+^, CD16^−^ and CD161^+^.^28, 29^ It has been reported to have anti-tumor activity in a broad spectrum of cancer cells through the perforin-granzyme pathway and secretion of Fas ligand and TNF-related apoptosis-inducing ligand.^29–31^We first confirmed whether this cell line showed any cytotoxic effects on the MDA-MB-231 breast cancer line we used through apoptosis induction. For that, we constructed a lentiviral FlipGFP vector (**Figure S2**, **Supporting Information**), which could allow us to detect cancer cell apoptosis directly without any additional staining, to distinguish fluorescence signals from T cells itself. In the FlipGFP structure, a caspase 3-cleavable peptide linker, DEVD, is added in GFP’s β-strand 11 to make it parallel to the β-strand 10, which deactivates the protein’s fluorescence due to the conformation mismatch. Activated caspase 3-triggered DEVD cleavage restores the antiparallel structure of β-strands 10 and 11, thereby reactivating the green fluorescence, so FlipGFP’s green fluorescence can represent a cell’s apoptotic activity.^32^ In addition to FlipGFP, our lentiviral vector encoded a constitutively expressed mCherry fluorescence protein, which allowed us to track the transduced cancer cells. We treated the FlipGFP-transduced MDA-MB-231 with staurosporine, a known apoptosis inducer,^33, 34^ and stained it with commercially available DEVD-linked blue fluorescent NucView® 405. At 5 h post-treatment, we observed that the FlipGFP signals (**Figure S3A**, **Supporting Information**) overlapped with the blue caspase-3 substrate signals (**Figure S3B**, **Supporting Information**) in these transduced (mCherry^+^) cells (**Figures S3C** and **S3D**, **Supporting Information**), confirming TALL-104’s cytotoxic effect on MDA-MB-231.

When incubated with TALL-104, MDA-MB-231 showed apoptosis signals (FlipGFP^+^) increasing with a higher the effector (TALL-104) to target (MDA-MB 231) ratio (E:T), indicating the cytotoxic effect of TALL-104 on MDA-MB-231 (**Figure S4A, Supporting Information**). We further confirmed TALL-104’s specificity on stromal cells using the same NucView® 405 substrate. Cancer cells (MDA-MB-231), endothelial cells (human microvascular endothelial cell; HMVEC) or normal fibroblast was incubated with TALL-104 in an E:T ratio of 2:1 for 2 h. By normalizing the caspase 3 substrate signal to that in absence of T cells for each condition, we found significantly higher caspase 3 activity in MDA-MB-231 compared against stromal cells (HMVEC and normal fibroblast; **Figure S4B, Supporting Information**). These results demonstrate that TALL-104 T cell-induced apoptosis is heavily weighted toward cancer cells, which is consistent with the report that TALL-104 can spare normal cells while killing cancer cells.^29, 30^ In addition, caspase-3 activity measured by the fluorogenic substrate matched the FlipGFP result.

We then used our recirculating tumor chip system to investigate the T cell infiltration in different cancer/stromal cell coculture conditions, with normal fibroblast (NF), cancer-associated fibroblast (CAF) and endothelial cell (EC). GFP-expressing MDA-MB-231 (termed 231), fibroblast (NF or CAF) and EC were seeded and cultured in our chip. After a two-day culture, CellTracker CM-DiI stained T cells (termed T; red) were introduced into the circulation (**Figure 3A**). The orthogonal projection of Z-stacked images obtained from the bottom chambers (**Figures 3B-3F**) clearly showed the T cell penetration to the tumor site (bottom chamber). For quantitative analysis, summation of the red signal from each Z-slice was normalized to the control signal without T cells for each group (**Figure 3G**). Compared with the coculture group with MDA-MB-231 and normal fibroblasts with endothelial cells (231/NF+EC+T), the absence of EC (231/NF+T) showed a significant T cell (red) signal increase in the bottom chamber, indicating that the endothelial monolayer hampered T cell infiltration. Anti-human PD-L1 antibody treatment (231/NF+EC+T+antiPDL1) enhanced T cell’s infiltration and even overcame the interference from endothelial monolayer, resulting in the highest T cell (red) signal amongst all the co-culture conditions with normal fibroblasts. PD-L1 is an immune checkpoint molecule that binds to its receptor (PD-1) on the T cells and the ligation of PD-1 delivers an inhibitory signal for T cell activation. Upregulated expression of PD-L1 have been found in many tumor cells and considered as one main factor arming tumors to escape from immune surveillance.^35^ Besides cancer cells, other cellular components in the tumor microenvironment also express PD-L1, such as macrophages, dendritic cells, activated T cells, CAFs, as well as ECs.^36, 37^ Expression of PD-L1 on ECs have been reported to compromise CD8^+^ T cell activation and cytolytic activity *in vitro*.^38^ Blocking PD-L1 on ECs promotes interferon gamma secretion and cytolysis by CD8^+^ T cells. In addition, elevated PD-L1 expression on tumor ECs is associated to lower CD8^+^ T cell infiltration and poorer prognosis.^39^ Our observation on elevated TALL-104 T cell infiltration across an endothelial monolayer under anti-PDL1 treatment correlates with the results from these studies, confirming that the critical role of endothelium in regulating tumor’s immune response could be recapitulated in our recirculating tumor chip system.

**Figure 3.**
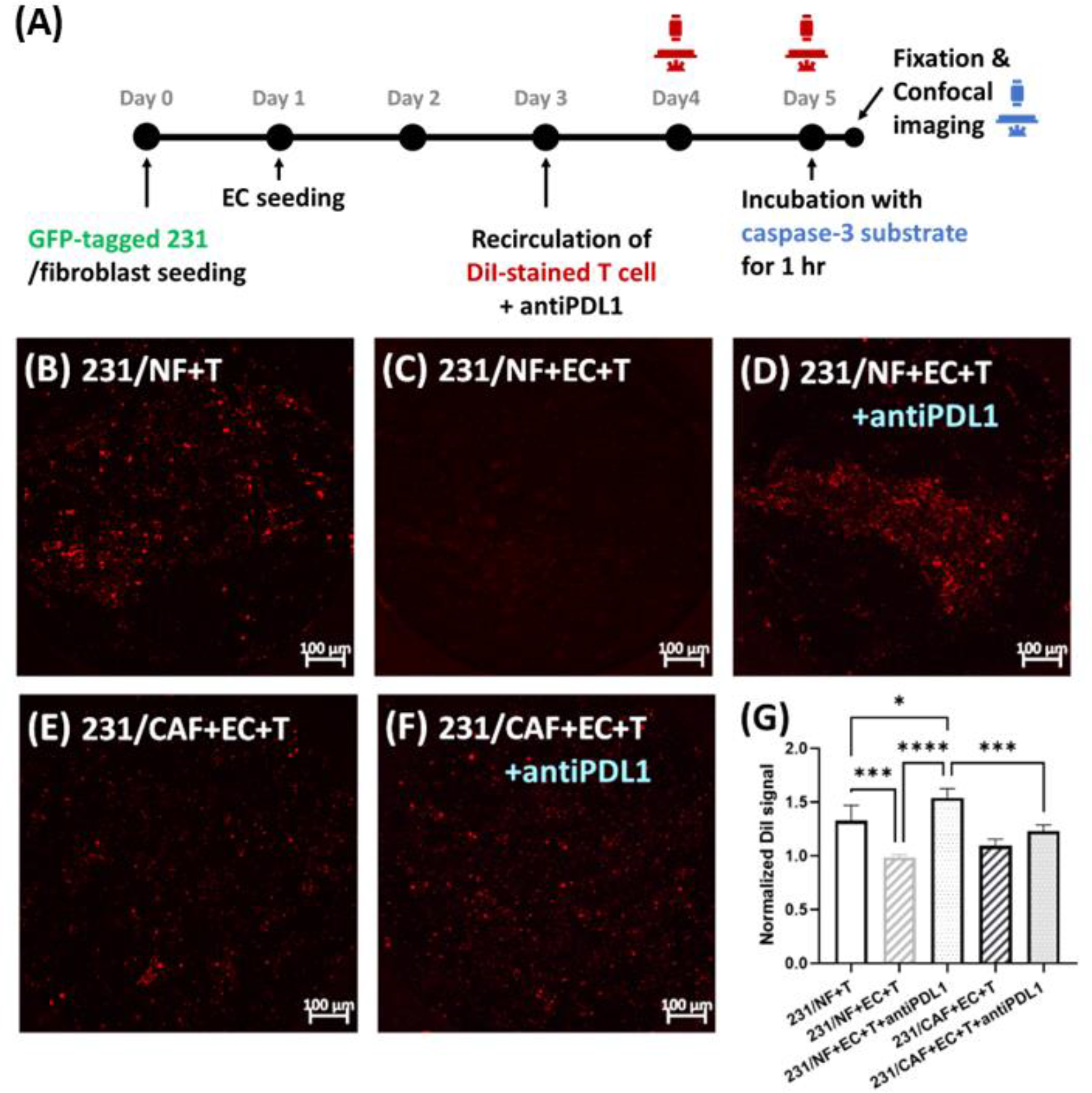
T cell infiltration in different coculture conditions. (A) Experimental design for studying of T cell infiltration in our system. Representative orthogonal projection images of DiI stained T cells in bottom chamber with coculture of (B) MDA-MB-231 and normal fibroblast (NF), (C) in the presence of endothelial cells, (D) with the treatment of anti-PD-L1 antibody, (E) in the coculture of MDA-MB-231, CAFs and endothelial cells, and (F) with the treatment of anti-PD-L1 antibody. (G) Quantitative analysis of the amount of DiI-stained T cell signals in the bottom chamber after 2-day recirculation. Significance was determined using one-way ANOVA with Tukey’s Multiple Comparison Test and presented as *, *p* < 0.05; **, *p* < 0.01, ***, *p* < 0.001; ****, *p* < 0.0001.

Compared with the coculture using normal fibroblasts, coculture with CAFs slowed down the T cell infiltration (**Figure 3G**). Studies have shown that CAF impedes T cell recruitment via multiple mechanisms.^40, 41^ For instance, CAF mediated ECM remodeling, which consequently suppressed the immune cell infiltration.^40, 42, 43^ In addition, depletion of FAP^+^ CAF increased CD8^+^ T cell recruitment in a murine TNBC model.^44^ In the other pancreatic ductal adenocarcinoma model, FAP^+^ CAF-derived CXCL12 was associated with T cell exclusion, and scavenging CXCL12 enhanced the presence of T cells in tumor.^45^ Similarly, knockdown of chitinase 3-like 1 (Chi3L1), also known as YKL-40 in human, in CAFs attenuated tumor growth and enhanced CD8^+^ T cell accumulation in breast tumor-bearing mice.^46^ In experiments using our system, we also found suppression of CD8^+^ T cell infiltration in the CAF-cocultured tumor microenvironment, drawing a correlation between our ex vivo findings and other *in vivo* results as well as demonstrating the feasibility of using our system to recapitulate key physiological conditions. The molecular mechanisms behind the CAF effects on T cell infiltration warrant further studies.

Caspase 3 activity of the tumor site (bottom chamber) on our chip was analyzed using the fluorogenic substrate to evaluate killing effects of recirculating T cells in all on-chip cocultured conditions shown in **Figure 3A**. As shown in **Figure 4A**, the caspase 3 activity observed in all NF-coculture groups was higher than the CAF-coculture groups. Although T cell alone did not induce more apoptosis, additional treatment with anti-human PD-L1 antibody boosted the cytotoxic effects in both NF- and CAF- coculture conditions. Notably, cytotoxic enhancement by anti-PD-L1 antibody in CAF-coculture group (231/CAF+EC+T+antiPDL1) was not able to achieve the level found in its NF counterpart (231/NF+EC+T+antiPDL1). The results reaffirm the crucial role of CAFs on fostering cancer cell survival and suggest that a combinational regimen of CAF normalization/depletion and anti-PD-L1 antibody therapeutics may lead to better therapeutic efficacy. Indeed, TGF-β secreted by CAFs^47^ has been reported to downregulate cytolytic protein expression in the CD8^+^ T cell, including perforin, granzyme, Fas ligand and interferon-γ, thereby compromising T cell’s cytotoxic activity.^48^ Moreover, melanoma-associated fibroblasts have been shown to decrease the NKG2D-dependent cytotoxic activity of natural killer cells by inhibiting the surface NKG2D ligand expression in tumor cells through matrix metalloproteinase secretion,^49^ which is another cancer cell-killing mechanism employed by TALL-104.^29^. Our result showing lower caspase 3 activity in tumor microenvironment with CAFs reflects this mediatory role of CAFs in immunosuppression.

**Figure 4.**
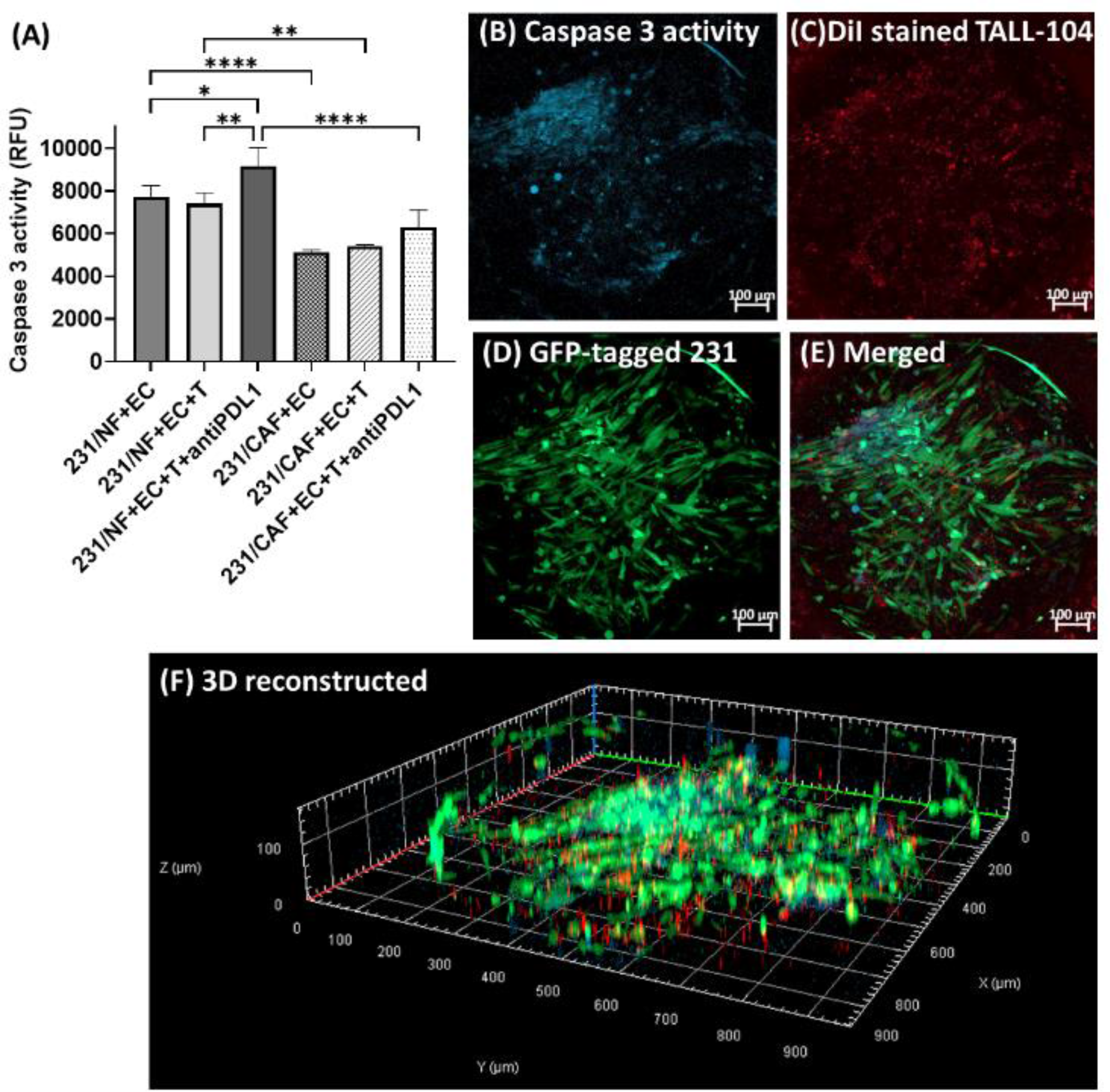
Apoptosis analysis in different TME conditions. (A) Quantitative analysis of T cell-induced apoptosis signals in different TME conditions. Results are presented as average ± S.D. (n = 3). Significance was determined using one-way ANOVA with Tukey’s Multiple Comparison Test and presented as *, p < 0.05; **, p < 0.01; ****, p < 0.0001. Representative images of the bottom chamber with the TME composed of MDA-MB-231, normal fibroblast, endothelial cell and TALL-104 with anti-human PDL1 antibody (the 231/NF+EC+T+antiPDL1 group). Maximum intensity projection of the full z-stack images captured from bottom to top layers for (B) caspase-3 activity, (C) DiI-stained TALL-104, (D) GFP-tagged MDA-MB-231 and (E) the merged image. (F) Representative 3D reconstructed image.

**Figures 4B** to **4F** are representative images of cancer cells cocultured with normal fibroblasts and ECs while CD8^+^ T cells and anti-human PD-L1 antibodies recirculated in the μFCA system. The blue signals from caspase 3 substrate mainly overlapped with green signals from GFP-expressing MDA-MB-231, indicating that the detected caspase 3 activity primarily came from cancer cells (**Figure 4B**). In addition, we observed that the infiltrated T cells (red signals) predominantly accumulated surrounding the cancer cells (green signals; **Figure 4C**). Studies have shown that cytotoxic T cells rearrange their cytoskeleton and secretory apparatus towards the point of contact with target cell during their interactions.^50–52^ The polarization and exocytosis of lytic granules then contribute to T cell’s cytotoxic effects. Our observation of T cells gathering towards the cancer cells in bottom chamber of the μFCA could highlight one of the key initiating steps of the contact-driven cytolytic process. Treatment of anti-PD-L1 antibody boosted the CD8^+^ T cell infiltration in the tumor microenvironment (**Figure 3**), and this could lead to enhanced downstream cytotoxic activities effected via more direct contacts between T cells and cancer cells (**Figure 4**).

In summary, by integrating new recirculating circuit with reconfigurable μFCA module designs, we developed a tumor-on-a-chip system that recapitulated the interactive dynamic interplay between immunotherapeutic agents (anti-PD-LI antibody), TME (MDA-MB-231, and stromal cells) and the immune cells (TALL-104 CD8^+^ T cell). The recirculating circuit enabled T cell circulation without compromising its viability, while the μFCA offered two different configurations (parallel or orthogonal), where the throughput can be scaled up from 8 to 8×8 conditions via orthogonally oriented bonding of the top and bottom layers of a μFCA and integration of pneumatic valves. Our results showed negative effects of stromal cells (endothelium and fibroblasts) on T cell infiltration. Both endothelium and CAFs could act to obstruct T cell infiltration, while anti-human PD-L1 antibody treatment could improve T cell infiltration. These results clearly showed the influence of tumor stroma to CD8^+^ T cell infiltration and its cytotoxic activity, corroborating with *in vivo* studies. Collectively, we demonstrated the feasibility of using our new recirculating tumor-chip system to study the cell-cell as well as cell-drug interaction dynamics, which can produce a better preclinical screening platform for the development of cancer immunotherapeutic agents.

## ASSOCIATED CONTENT

### Supporting Information

Materials and methods section, Figures S1-S4: Representative micrograph-derived images and leakage validation of the pneumatic valve we designed; plasmid map of lentiviral FlipGFP; FlipGFP-response to caspase 3 stimulation in the FlipGFP-transduced MDA-MB-231; cytotoxic effect of TALL-104 to MDA-MB-231.

## ACKNOWLEDGMENT

The authors acknowledge the instrumental support from the nanofabrication facility at CUNY-Advanced Science Research Center. This work is supported by Pershing Square Sohn Cancer Research Alliance (S.W.), American Society of Cell & Gene Therapy (Career Development Award, ASGCT2021003, Y.-H.L.) and NIH (UG3/UH3TR002151, K.W.L.).

