## Supporting Information for "An Immune Cell Recirculation-Enabled Microfluidic Array to Study Dynamic Immunotherapeutic Activity in Recapitulated Tumor Microenvironment"

**This document includes**

Materials and Methods
Figure S1. Representative micrograph-derived images and leakage validation of the pneumatic valve we designed
Figure S2. Plasmid map of the lentiviral FlipGFP vector constructed in this study.
Figure S3. FlipGFP-response to caspase 3 stimulation in the FlipGFP-transduced MDA-MB-231.
Figure S4. Cytotoxic effect of TALL-104 to MDA-MB-231.

**Materials and Methods**

*Microfluidic device fabrication*. We followed our protocol in a previously published work to fabricate the device with minor modifications.[1, 2] Briefly, the SU8 master molds were made on silicon wafer substrates by photolithography, to 100µm and 200µm heights for the top channel and the bottom chamber layers in our microfluidic cell array (µFCA), respectively. For the recirculating circuit, the height of SU8 molds were constructed to a height of 200µm for both its top and bottom layers. SU8 pillars of 70 µm height served to create a pattern of holes across the thin middle (membrane) layer. Polydimethylsiloxane (PDMS, DOW Chemical, Midland, MI) was spin-coated and cured on the middle master for the thin membrane. PDMS was directly cast on the top and bottom layer masters. Bonding between layers was enabled after oxygen plasma treatment, aligning layers with the aid of a stereomicroscope (Motic Scientific, Schertz, TX). Selective bonding was done by transferring PDMS residual oligomers from a PDMS stamp to the stopper region of oxidized (plasma treated) bottom layer before bonding with the bonded top and middle layers.[3] **Figures 2B** and **2C** in the main text show the design of our high-throughput µFCA configuration with pneumatic valves. The opening pore area ratio in the middle layer was unchanged from the original 8 condition design, to maintain the same permeability of endothelium. The normally closed valve comprised a control channel in the top layer, a stopper in between two adjacent bottom chambers and a deformable middle layer, which was not bonded to the stopper. It allowed flow to go through the bottom channel while a negative pressure on the control channel was applied.

*Cell culture*. Human breast cancer cell line MDA-MB-231 (HTB-26TM) and human normal fibroblast CCD-1096Sk (CRL-2129) were purchased from American Type Culture Collection (Manassas, VA). MDA-MB-231 cells and normal fibroblasts were cultured in high glucose

DMEM (Thermo Fisher, Waltham, MA), supplemented with 10% Fetal Bovine Serum (FBS; Atlanta Biologicals, Atlanta, GA) and 1× penicillin-streptomycin (Thermo Fisher). Human dermal blood microvascular endothelial cells (HMVECs) were cultured in the EGM™-2 basal medium with growth factor supplements (Lonza, Switzerland). Human mesenchymal stem cells (MSCs) were cultured in low glucose DMEM (Thermo Fisher) supplemented with 10% FBS and 1× penicillin-streptomycin, and the seeding density of each subculture was 5000 cells/$cm^2$. Cancer-associated fibroblasts were differentiated from human MSCs by culturing MSCs in the MDA-MB-231-conditioned media for 30 days by following the protocol from previously published works.[2, 4, 5] Human monocyte line, THP-1(TIB-202™) from ATCC, was cultured in RPMI-1640 (Thermo Fisher) supplemented with 10% FBS and 0.05mM β-mercaptoethanol. Primary mouse CD8+ T cells were maintained in RPMI-1640 supplemented with 10% FBS, 1× nonessential amino acids, 2 mM L-glutamine, 1 mM sodium pyruvate and 50 μM β-mercaptoethanol. A human $CD8^+$ T cell line, TALL-104 (CRL-11386) from ATCC, was cultured in RPMI-1640 supplemented with 20% FBS and 100 units/mL IL-2 (Miltenyi, Germany).

*3D coculture in the tumor-chip system*. The device was initially UV sterilized in a cell culture hood for at least 1h. In order to enhance cell attachment, fibronectin bovine plasma (30 μg/mL; Sigma-Aldrich, St. Louis, MO) was introduced into the device and incubated at 37°C for at least 1h. The device was then filled with the complete medium. After this pretreatment, the device was stored at 4°C. Cancer cells or the mixture of cancer cells and stromal cells (1:1) at a concentration of $2\times10^7$ cells/mL were mixed with 4 mg/mL growth factor-reduced Matrigel (Corning Inc., Corning, NY), and the cell-containing Matrigel was subsequently loaded into the bottom chambers. Inlets and outlets of the top channel were clamped shut during the cell seeding process, so that there was no pressure gradient to allow pressure-driven flow between the top channel and the bottom chambers, effectively preventing crosstalk. After gelation at 37$^{o}$C for 1h, the clamped top channel was released and, reversely, inlets and outlets of the bottom chamber were clamped closed. Fresh medium was then continuously introduced into the top channel using a syringe pump at a speed of 100 µm/s. The device was incubated at 37$^{o}$C overnight under 5% $CO_2$ atmosphere. Afterwards, HMVECs were seeded into the top channel at a density of $1\times10^7$ cells/mL. Excess HMVECs were removed at 1h post-seeding. Again, the device was incubated at 37$^{o}$C and under 5% $CO_2$ atmosphere; continuous medium flow was given after HMVECs attached to the channel (at ~6 h post-seeding). A combinational medium composed of EGM$^{TM}$-2MV and DMEM in a ratio of 1:1 was used in all of our culture conditions.

*Establishment of FlipGFP-expressing MDA-MB-231.* The lentiviral vector encoding FlipGFP-T2A-mCherry was first constructed by PCR cloning. Briefly, the FlipGFP-T2A-mCherry sequence was first retrieved from the original FlipGFP plasmid[6] (Addgene #124428) using the Thermo SuperFi high-fidelity polymerase with the following specific primers:

forward: TCGTGACGTACGCCACCATGGACCTGCCTGA;

reverse: GATATCGAATTCTTACTTGTACAGCTCGTCCATGCC. The Lentivral backbone (Addgene #61422, Watertown MA) and the PCR product were then digested by the restriction enzymes, BsiWI and EcoRI (NEB, Ipswich MA). After purification, the backbone and insert were ligated using the T7 ligase at 25$^{o}$C for 30 mins. The ligated product was transformed into the Takara Stellar competent cell (Japan) and purified using the NucleoBond® Xtra Midi Plus Endotoxin-Free kit. The final Lentiviral FlipGFP vector sequence was verified by Sanger sequencing (Genewiz, South Plainfield NJ), and its function was confirmed by in vitro transfection on the HEK293T cell. For Lentivirus production, the Lentiviral FlipGFP vector was co-transfected to the HEK293T cell with the packaging plasmids (psPAX2 and pMD2.G, Addgene) using Calfectin (SignaGen Laboratories, Frederick MD) by following manufacturer's instruction. The virus-containing media were collected at 48h and 72h post-transfection. Afterwards, viruses were purified using a 0.45 µm cellulose filter (VWR, Radnor PA) and concentrated using AMICON 100K MWCO column (Millipore-Sigma). Virus titer was then quantified by RT-qPCR. The breast cancer cell line (MDA-MB-231) was transduced with the produced viruses with MOIs of 5, 10 and 20 in the presence of polybrene (8µg/mL, Millipore-Sigma), and the $mCherry^+$ cells were then sorted at 48h post-transduction for further expansion.

*TALL-104 cytotoxicity in the 2D assay*. MDA-MB-231, HMVECs or human normal fibroblasts were seeded in a 48-well plate with a density of 40,000 cells per well. Biotium NucView®405 Caspase-3 enzyme substrate (Fremont, CA) was used for monitoring the cytotoxicity of $CD8^+$ T cells. TALL-104 was added to the plate with the effector cell-to-target cell ratios (E:T) of 1:1, 2:1, 5:1 and 10:1, along with 2µM caspase-3 substrate. Cells were then imaged at 2h post-treatment using the fluorescent microscope (Zeiss AxioObserver)

*$CD8^+$ T cells infiltration and cytotoxicity in µFCA*. The general experimental timeline is presented in **Figure 3A** of our main text. GFP-expressed MDA-MB-231 was first mixed with normal fibroblast or CAF in Matrigel and loaded into the bottom chambers as aforementioned. Endothelial cells were then introduced on the second day, and a continuous flow of media was given at 5 h post-seeding. At Day 4, TALL-104 was first stained with 2 µM CellTracker CM-DiI (Thermo Fisher) and washed with PBS by following manufacturer's instruction. Prior to T cell introduction, the recirculating circuit was connected to the µFCA. Anti-human PDL1 antibody (Bio X cell, Lebanon, NH) was mixed with the T cell suspension in the medium. The syringe pump was set to

switch direction every 300 µl infusion. The Z-stacked images for monitoring T cell infiltration were captured by the fluorescent microscope (Zeiss AxioObserver, Germany) each day after T cell introduction. At the endpoint, cells were incubated with 2µM NucView®405 caspase-3 substrate for 1 h. Cells in the device were then fixed with 4% paraformaldehyde for 60 min. To avoid the interference of emission by GFP-expressed MDA-MB-231, caspase-3 activity was measured using a confocal microscope with Z-stacked sectioning (Zeiss LSM800).

### Supplementary Figures

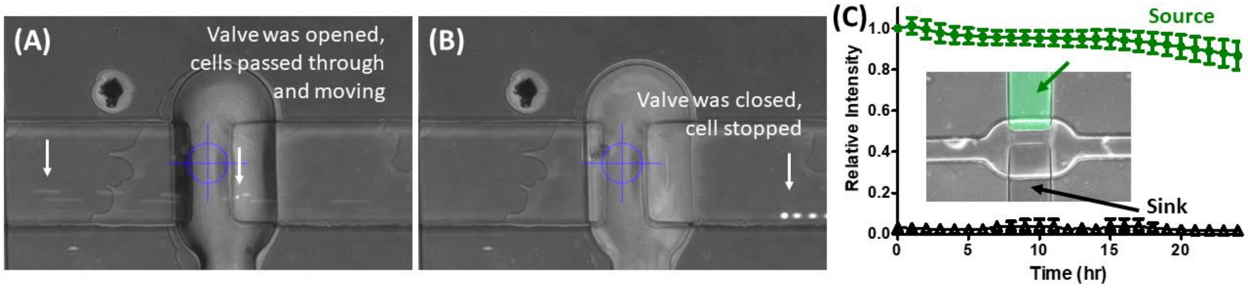


**Figure S1.** Representative micrograph-derived images and leakage validation of the pneumatic valve we designed (A) Representative image showing that CellTracker-stained fibroblasts passed through the valve when the valve was opened. The image frames from a 2-minute movie were extracted and stacked together for image generation. (B) Representative image showing that the fibroblasts stopped when the valve was closed. (C) Relative intensity changes of sodium fluorescein in the source and receiver chambers over time. Results are presented as average ± standard deviation (S.D.; n = 3).

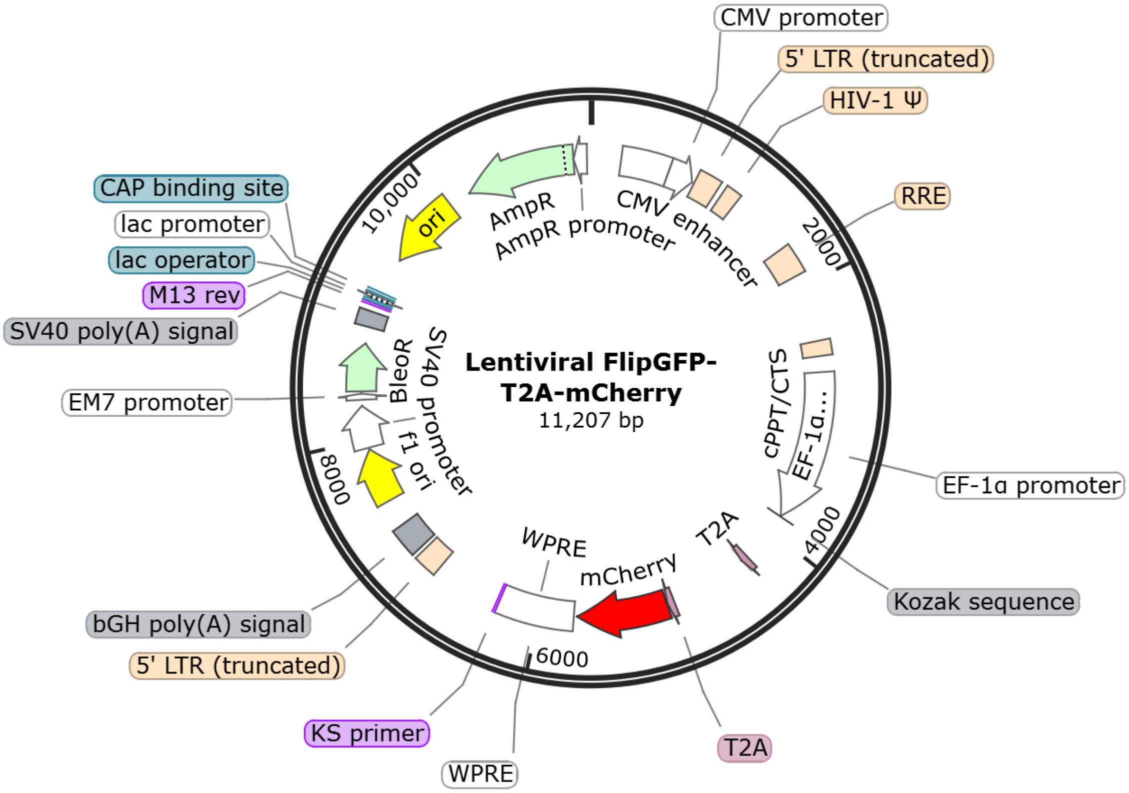


**Figure S2** Plasmid map of the lentiviral FlipGFP vector constructed in this study.

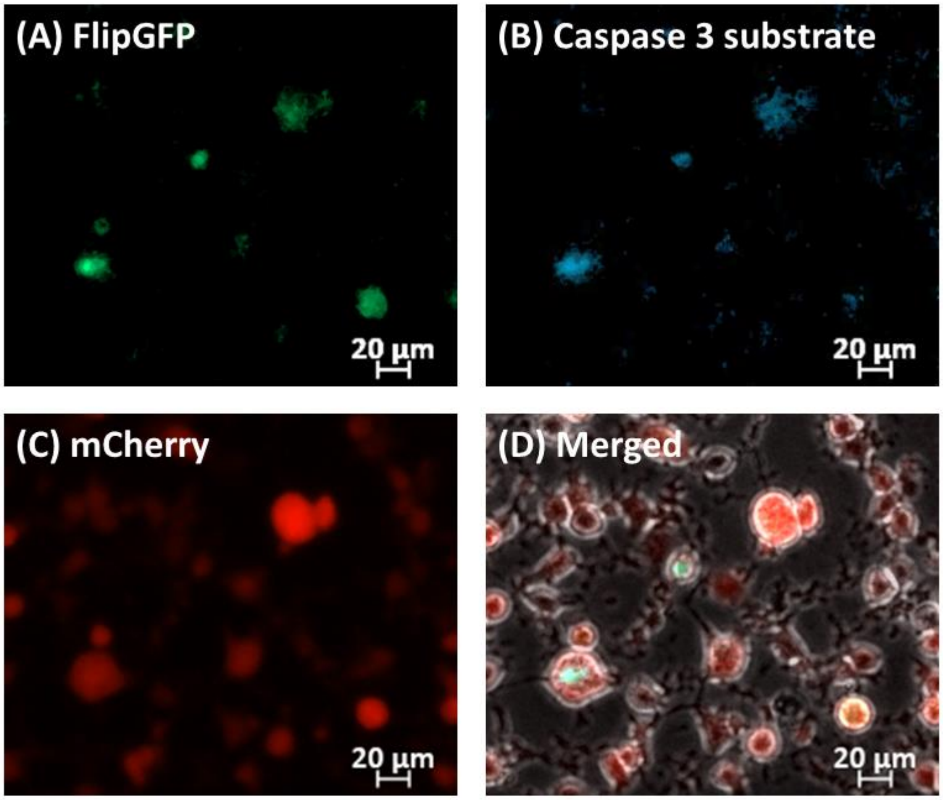


**Figure S3.** FlipGFP-response to caspase 3 stimulation in the FlipGFP-transduced MDA-MB-231. (A) FlipGFP signals and (B) caspase 3 substrate signals from the FlipGFP-trasnduced MDA-MB-231 after being treated with 1µM staurosporine treatment for 5h. (C) mCherry signals from the transduced MDA-MB-231 cells. (D) Merged image of the phase contrast with all fluorescent channel (B-C; green, blue, and red) images to confirm the overlap between FlipGFP and caspase 3 substrate signals.

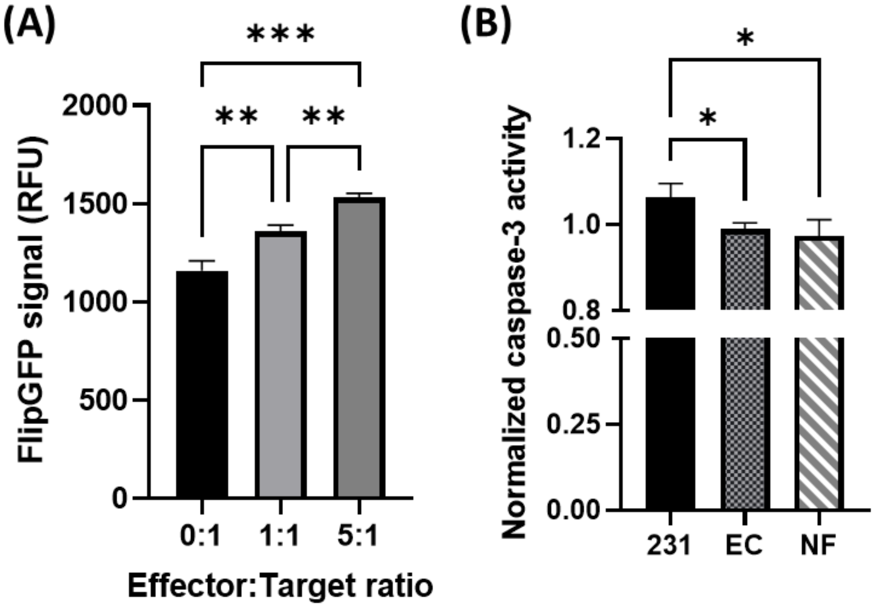


**Figure S4.** Cytotoxic effect of TALL-104 to MDA-MB-231 and stromal cells. (A) FlipGFP signal quantification from the MDA-MB-231 cells incubated with TALL-104 in different effector: target (E:T) ratios for 3h. (B) TALL-104 cytotoxic effect on different cell types (cancer cells, endothelial cells or normal fibroblasts). Cells were incubated with TALL-104 in an E:T ratio of 2:1 for 2h and then stained with the caspase-3 substrate. Results are presented as average ± S.D. (n = 3). Significance was determined using one-way ANOVA with Tukey's Multiple Comparison Test and presented as *, $p < 0.05$; **, $p < 0.01$, ***, $p < 0.001$.
